# Automated Reactive Accelerated Aging for Rapid *In Vitro* Evaluation of Neural Implants Performance

**DOI:** 10.1101/204099

**Authors:** Matthew G. Street, Cristin G. Welle, Pavel A. Takmakov

## Abstract

**Objective:** Novel therapeutic applications for neural implants require miniaturized devices. Pilot clinical studies suggest that rapid failure of the miniaturized neural implants in the body presents a major challenge for this type of technology. Miniaturization imposes stricter requirements for reliability of materials and designs. Evaluation of neural implant performance over clinically relevant timescales presents time-and cost-prohibitive challenges for animal models.

**Approach:** *In vitro* reactive accelerated aging (RAA) was developed to expedite durability testing of these devices. RAA simulates an aggressive physiological environment associated with an immune response and implicated in device failure. It uses hydrogen peroxide, which mimics reactive oxygen species (ROS), and high temperature to accelerate chemical reactions that lead to device degradation. RAA accurately simulates the degradation pattern of neural implants observed *in vivo*, but requires daily maintenance and is prone to variability in performance.

**Main results:** This work introduces automated reactive accelerated aging (aRAA) that is compatible with multiplexing. The core of aRAA is electrochemical detection for feedback control of hydrogen peroxide concentration, implemented with simple off-the shelf components.

**Significance:** aRAA allows multiple parallel experiments for a high-throughput optimization of reactive aging conditions to more quickly and more rigorously simulate the *in vivo* environment. aRAA is a cost-effective tool for rapid *in vitro* evaluation of durability of neural implants, ultimately expediting the development of a new generation of miniaturized devices with long functional lifespans.

## Introduction

Neural implants are medical devices that modulate and record nervous system activity for therapeutic applications. Common clinical examples of implantable neurological devices are cochlear implants for restoration of hearing, spinal cord stimulators for pain relief and deep brain stimulators for treatment of tremor (Kumsa et al., 2017). The repertoire of diseases to be addressed with neural implants is continuously expanding with the increased attention on innovative applications (Birmingham et al., 2014; Miranda et al., 2015). Cortical neural implants for brain-computer interfaces (BCIs) have evolved from a passion of a small group of scientists (Gay, 2015) into a family of clinical devices with defined roadmaps to clinical use (Bowsher et al., 2016). Advancements of peripheral nerve stimulators drive bioelectronic medicine, where targeted neuromodulation of nerves connecting peripheral organs is used to treat a variety of medical conditions with unprecedented flexibility and potential to mitigate the side effects (Waltz, 2016).

Novel applications for neural implants require higher spatial resolution to increase “bandwidth” of the communication between medical devices and a nervous system. Progress in neuroprosthetic BCI development is measured by the number of neurons that can be recorded simultaneously (Stevenson and Kording, 2011), which requires microelectrodes of dimensions comparable to a size of a single neuron (< 100 micrometers). Additionally, use of sensory feedback for a neuroprosthetic relies on high spatial resolution of electrical stimulation of neurons responsible for tactile sensation. Furthermore, a favorite target for bioelectronic medicine modulation is the vagus nerve, which is a nervous system “super-highway” that is 100,000 axons wide, where spatially focused stimulation might be the key to eliminating stimulation-induced side effects.

An increase in spatial resolution requires miniaturization of neural implants, which is accomplished using microfabrication methods developed in the microelectronic industry. However, miniaturization of these devices brings additional challenges associated with rapid decline in device performance that has been extensively documented and considered to be a great challenge by the neuroengineering community (Takmakov, 2017; Wellman et al., 2017). Discovery of new materials and development of new device designs for a reliable miniaturized neural interface requires rapid and high-throughput reliability testing methods. However, the performance testing for traditional clinical neural implants has typically been done in large animals (Kumsa et al., 2017) due to the large size of the implants and to ensure clinical translation of the study results (Hachmann et al., 2013). While miniaturization of neural implants enabled testing of their performance in rodents (Vasudevan et al., 2017), all animal experiments are expensive, lengthy, require highly-skilled personnel and subject to a variability. To expedite neural implant development, we designed a reactive accelerated aging (RAA) (Takmakov et al., 2015) for rapid simulation of in vivo degradation of these devices using hydrogen peroxide (H_2_O_2_) (Patrick et al., 2011), that mimicked reactive oxygen species (ROS) associated with immune system attack (Potter-Baker and Capadona, 2015) and high temperature (Hukins et al., 2008) to accelerate the chemical reactions. The RAA provided valuable information on failure modes of cortical neural implants that included metal dissolution, moisture penetration, degradation and delamination of insulation. The patterns of neural implant degradation observed after 7 days in RAA appeared very similar to data from chronic animal studies (Barrese et al., 2013, 2013; Prasad and Sanchez, 2012; Prasad et al., 2014). Furthermore, a recent analysis of data on explanted cochlear implants pointed to importance of ROS in degradation of insulation and electrodes (O’Malley et al., 2017), suggesting that RAA accurately simulates the in vivo environment.

In a short time, RAA became a very popular testing platform with many researchers replicating the design (Caldwell et al., 2018; Pfau et al., 2017) due to its ability to provide data on materials reliability and robustness of the implant design without the need for animal experiments. However, the original RAA design (Takmakov et al., 2015) was complex and required constant human attention, which prevented wider adaptation of the RAA. We have gone through a several iterations in developing an automated RAA, incorporating optical and electrochemical detection and have devised what we now present as a robust and reproducible design for the automated RAA (aRAA) that is simple to build and easy to operate. The key part of automation is a feedback loop that relies on electrochemical activity of H_2_O_2_

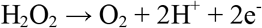

This method of H_2_O_2_ measurement is ubiquitously used in enzyme-coupled biosensors including clinical devices suggesting that it is a simple and robust analytical technique. We have developed chronoamperometric (constant potential pulses) detection of H_2_O_2_ concentration on platinum microelectrodes, which enable accurate and stable measurement of H_2_O_2_ over a long period of time (weeks). Additionally, we have modified the system so it can be built using off-the-shelf Arduino and Raspberry Pi based parts. The new aRAA design allows easy scaling of the RAA setup to have multiple modules running in parallel at different conditions (H_2_O_2_ concentration, temperature and time) to precisely tune the intensity of reactive aging. To take advantage of this new feature, we performed simultaneous reactive aging of Blackrock and TDT microelectrode arrays (MEAs) at two different temperatures (87° C and 67° C), characterizing the degradation of each using electron microscopy and impedance spectroscopy. Data on reactive aging-induced degradation of Blackrock confirmed that the aRAA design has the same performance as original the RAA system. Milder degradation of TDT at the lower temperature demonstrates that fine tuning of RAA conditions is possible to better match in vitro reactive aging conditions to *in vivo* environment, and can be easily implemented in a new aRAA design.

## Methods

### Overview of Automated Reactive Accelerated Aging System

The automated reactive accelerated (aRAA) system with two independent modules, each of which uses different sets of experimental parameters, was implemented (Figure 1). Each reaction vessel consisted of a five-neck European-style 125mL flask (Ace Glass Inc., Vineland, NJ), a PID temperature controller, an electric heating mantle, an electrochemical H_2_O_2_ feedback controller, a three-channel peristaltic pump (400DM3, 120s, Watson-Marlow, Wilmington, Massachusetts), and a magnetic stirrer (Nuova ii, Ramsey, Minnesota). Each reaction flask had two small ports dedicated to Pt working microelectrode and carbon rod counter electrode inserted with the stoppers that came with the flask. Left and right side ports were used to hold thermocouple and tubes for delivery of the liquids via 14/20 PTFE cap with an opening (Cat# F20309-1680, belart.com). Microelectrode arrays for testing were placed in a central port with Blackrock being held by a custom build PTFE holder and TDT array being taped with PTFE tape to a PTFE rod fixed in an opening of 24/40 PTFE cap (Cat# F20311-1718, belart.com).

**Figure 1.**
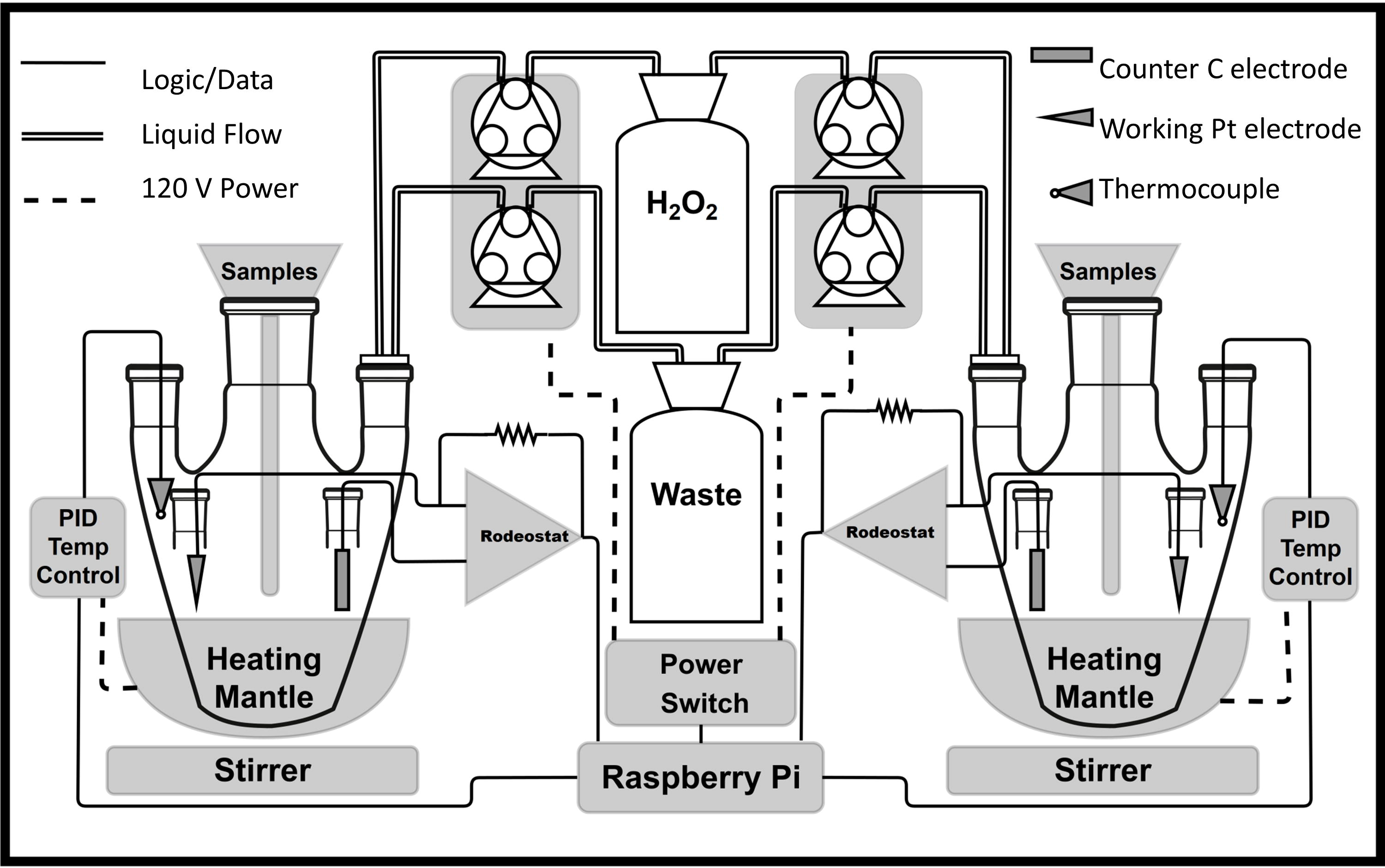
Diagram of automated reactive accelerated aging (aRAA) system with feedback loop for maintenance of H_2_O_2_ concentration. Each aRAA module is a five-neck flask with a magnetic stirrer and is equipped with feedback loops to maintain desired temperature and H_2_O_2_ concentration. Temperature feedback loop consists of heating mantle connected to PID temperature control unit. H_2_O_2_ concentration is measured electrochemically using Rodeostat, an Arduino-based potentiostat, in a two-electrode scheme with platinum microelectrode as a working electrode and carbon rod as a counter electrode. Temperature and H_2_O_2_ concentration is sampled by Raspberry Pi control unit that activates pumps via power switch to deliver stock H_2_O_2_ solution whenever H_2_O_2_ concentration falls below a threshold value and to remove waste to maintain constant volume.

Automation was implemented with a Raspberry Pi that activated the peristaltic pumps to deliver H_2_O_2_ as necessary and logged the temperature and H_2_O_2_ concentration for each reaction module. The pumps delivered concentrated H_2_O_2_ (1.5 M) and removed excess of a solution to maintain constant volume using ~ 16 cm pieces of soft Viton tubing (1/16” ID 1/8 OD, Cat# 5119K39) that was connected to PTFE tubing (1/16” ID 1/8 OD, Cat# 5239K24) using barbed connectors (Cat# 53055K111, McMaster-Carr, Robbinsville, NJ). The Raspberry Pi was equipped with a battery-backed PCF-8523 clock (Adafruit, New York City, NY) to account for possible system power loss and to facilitate precise data logging. A 5 port powered USB extension was added to 4 on board Raspberry Pi USB ports to enable connection to two Rodeostat potentiostats, two PID temperature controllers, a keyboard and a mouse. The main script running on the Raspberry Pi was written in Python with the purpose of maintaining serial communication with the peripherals to read and log process values as well as set them as necessary. Finally, the Raspberry Pi was configured to initialize the script at startup in the case of power failure and to run it until the specified end date for each module was reached. All solutions were prepared with deionized water (18MΩ cm) using PBS tablets and 30% H_2_O_2_ solution (Fisher Scientific, Hampton, NH). To initialize an experiment, flasks were filled with a H_2_O_2_ solution in PBS (15 mM).

### Maintenance of Temperature

Each reaction vessel was equipped with a hermetically sealed PFA jacketed thermistor (Omega Engineering, Inc., Norwalk, Connecticut) connected to a Platinum Series PID controller (Omega Engineering, Inc., Norwalk, Connecticut). PID parameters were tuned for each individual reaction module using the embedded auto-tune feature of the PID device. Each PID controller powered a DC-controlled solid state relay (Omega Engineering, Inc., Norwalk, Connecticut) using on/off control of a resistive element heating mantle (Glas-Col LLC, Terre Haute, Indiana). The PID controllers were connected to the Raspberry Pi via USB to set or record module temperature.

### Maintenance of Hydrogen Peroxide Concentration Using Electrochemical Detection

H_2_O_2_ concentration was measured using electrochemical detection with chronoamperometry. In this technique voltage pulses were applied to 25 μm platinum disk working electrode (Figure 2A) and current was measured in a two electrode scheme with a carbon rod of 0.25” diameter (Electron Microscopy Sciences, Fort Washington, Pennsylvania) as a counter electrode. The platinum microelectrode was fabricated in-lab using platinum wire and glass capillaries according to procedure (Caruana and Bannister, 1997) detailed in a section below. The electrochemical experiments were performed with Rodeostat (IO Rodeo, Pasadena, California). The Rodeostat is an Arduino-based open source potentiostat with established Python libraries making automated use simple through using a Raspberry Pi. The parameters for chronoamperometry were determined using cyclic voltammetry (300 samples per second, scan rate 500 mV/sec, 2-electrode scheme) by finding the lowest anodic voltage at which H_2_O_2_ oxidized (Figure 2B). Microelectrodes were used to ensure that hemispherical diffusion profile with a steady state current is rapidly established. Faradaic current from H_2_O_2_ oxidation was measured after discharge of a capacitance associated with a double layer (Figure 2C). Optimal chronoamperometric waveform for detection of hydrogen peroxide was to hold electrode at - 0.3 V for 0.5 second and 2 second + 0.7 V pulses (Figure 2D). The current was detected by acquiring 10 samples over 1 sec at the end of anodic pulses and was converted to H_2_O_2_ concentration using a calibration curve established for each individual Rodeostat and Pt microelectrode pair. Each Rodeostat was connected to the Raspberry Pi using USB serial communication and controlled via the main Python script. To account for the noise generated in the signal by the magnetic stir bar, a running average with 10 data points of the detected current was used to calculate H_2_O_2_ concentration. The peristaltic pump on/off state was determined using a simple thresholding technique by the Raspberry Pi. When toggled on, the pump would deliver concentrated H_2_O_2_ to the module while simultaneously removing solution. In three channel pump head, one channel was for the delivery of H_2_O_2_ while two tubes were for the waste removal. The waste removal tubing was positioned in the flask such that the tip was at the same level as the surface of solution at the target volume. This arrangement allowed the removal of liquid above the level of the exhaust tube; at a rate double that of the H_2_O_2_ delivery, thus ensuring that the solution level remained constant.

**Figure 2.**
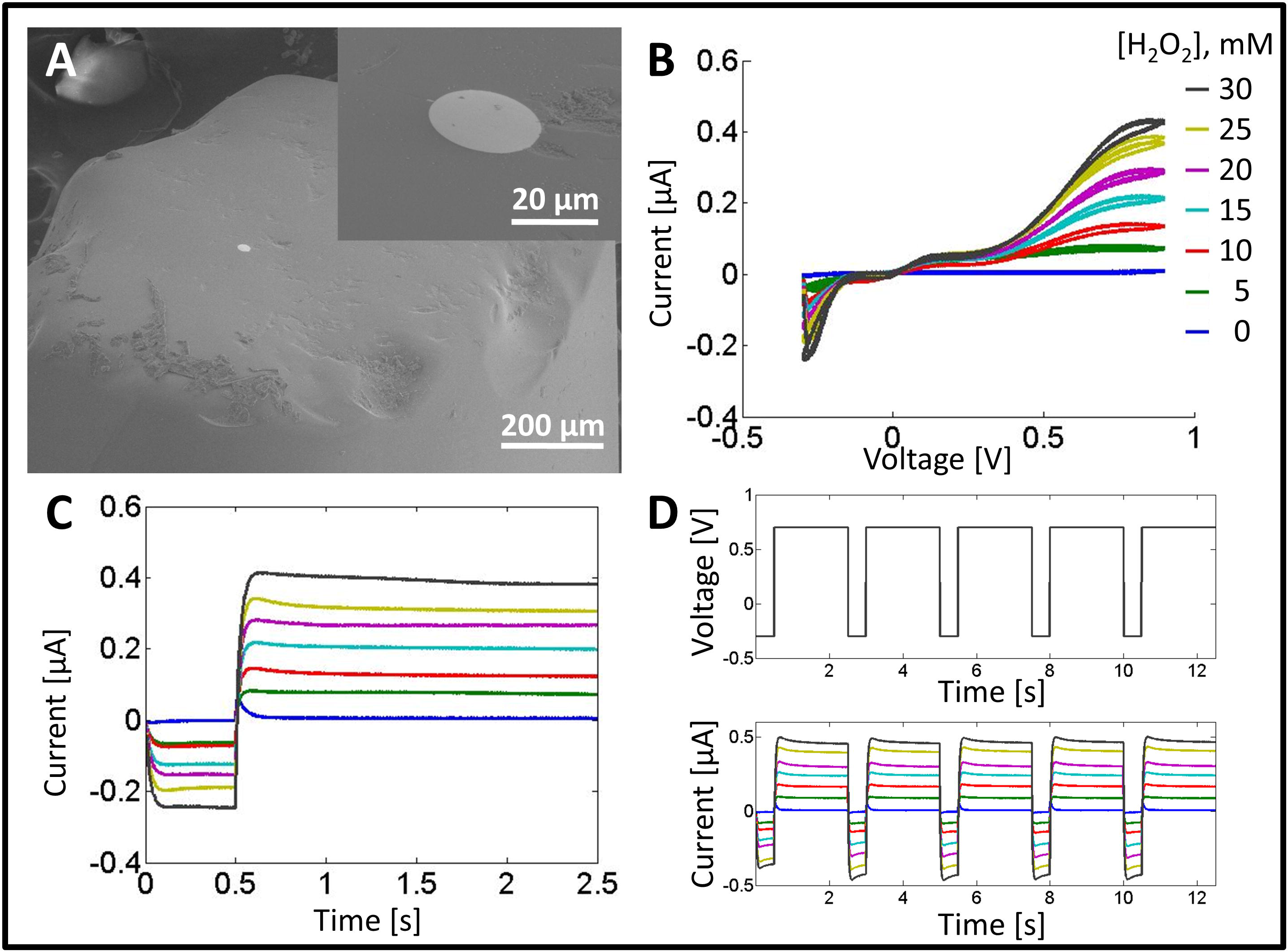
Electrochemical sensing for a feedback loop to maintain H_2_O_2_ concentration in the automated RAA. Detection of H_2_O_2_ is performed with 25 micrometer platinum disk microelectrodes (**A**) electron micrograph of the electrode with a higher magnification insert) using Rodeostat, an Arduino-based potentiostat. Optimum conditions for electrochemical detection were determined using cyclic voltammetry (**B**). The measurement of H_2_O_2_ concentration is done using chronoamperometry with 2 second anodic pulses to + 0.7 V being applied to microelectrode that is held at - 0.3V between the pulses (**C**). The current reached steady state very quickly which allowed for sampling every 3 seconds (**D**). Insert in (B) with color legend for 0 (navy blue), 5 mM (green), 10 mM (red), 15 mM (blue), 20 mM (purple), 25 mM (gold) and 30 mM (black) of hydrogen peroxide correspond to all plots with current traces.

### Fabrication of Platinum Microelectrodes

Platinum microelectrodes were fabricated in-house using 1.5/0.84 mm OD/ID glass capillary (1B150F-4, World Precision Instruments, Sarasota, FL); 25 μm platinum (Pt) wire (Alfa Aesar, Tewksbury, MA); hook up copper (Cu) wire and Sn63/Pb37 0.63 mm solder wire. First, one end of the glass capillary was sealed with a butane torch making the seal as small as possible. A 1.5 cm long piece of Pt wire was degreased in acetone for one minute and then inserted in the open end of the capillary. The sealed end of the capillary was pointed down and gently tapped on a hard surface until the Pt wire sled to the sealed bottom of the capillary. With the Pt wire at the sealed tip, the open end of the capillary was wrapped with 1-2 layers of parafilm and attached to a rubber tube of small diameter to create a good seal and then hooked to a vacuum pump. The capillary was fixed on a ring stand with the sealed end pointing downward at a 45° angle to the ground. The vacuum was turned on and the sealed end was carefully melted using butane torch. The glass was fused continuously for at least 2 mm from the tip of the Pt wire as to create a good seal around the Pt. After Pt wire was sealed, the working end of the capillary was ground with 600 grit sandpaper until the Pt wire is exposed. The microelectrode was then polished with a polishing kit first with 0.3 μm followed by 0.05 μm Al_2_O_3_ slurry (eDAQ, Australia). A 1 cm piece of solder was inserted in the open end of the capillary. The Cu wire was used to push the solder down the capillary until it reached the Pt wire. The capillary was slowly heated from the outside at the position of the solder using the butane torch. Once a good connection is formed between the Pt and Cu wire, the Cu wire was fixed to the capillary with a heat shrink as to remove mechanical stress from the soldered connection. The quality of a glass seal for Pt disk microelectrode was verified with electron microscopy (Mira3, Tescan USA Inc, Warrendale, PA) and via cyclic voltammetry of potassium ferrocyanide (Sur et al., 2012).

### Automated Reactive Accelerated Aging of Neural Implants

Commercial neural implants were subjected to RAA conditions to compare performance of aRAA system to its earlier version (Takmakov et al., 2015) using the same characterization protocol with electron microscopy and electrochemical impedance spectroscopy. In the first aRAA vessel, Blackrock Microsystems implant with 16 individual microfabricated electrodes coated with Parylene-C in 4 by 4 configuration (Blackrock Microsystems, Salt Lake City, UT) has been exposed to 15 mM H_2_O_2_ at 87 °C. In second aRAA module, TDT implant (TDT, Alachua, FL) with an array of 16 polyimide insulated gold-coated tungsten microelectrodes were exposed to 15 mM H_2_O_2_ at 67 °C. Experiments were run in parallel for 7 days with temperature and H_2_O_2_ concentration being logged with Raspberry Pi for both modules (Figure 3).

**Figure 3.**
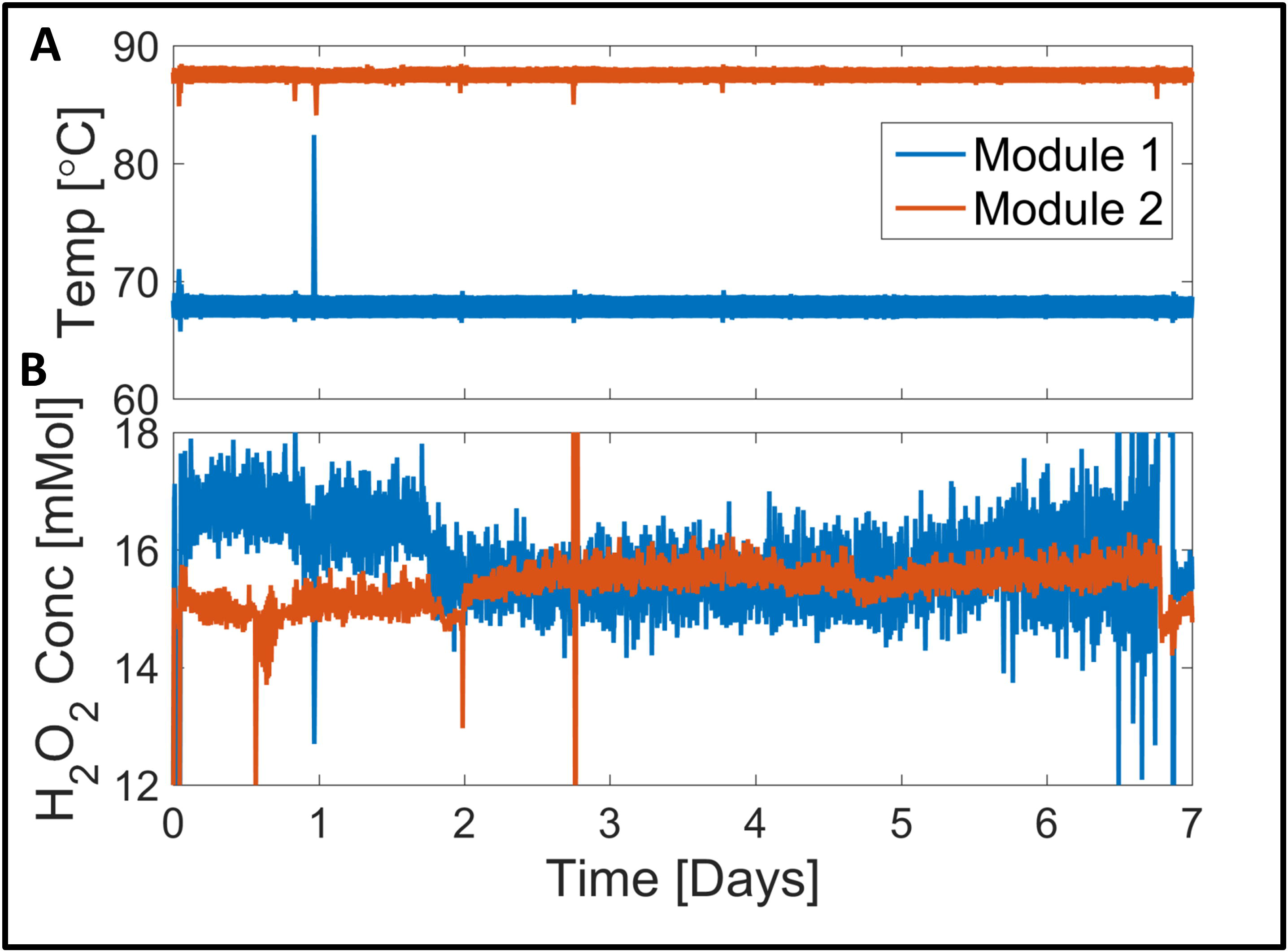
Performance of automated RAA system with two independently controlled modules. (A) Temperature and (B) H2O2 concentration for two RAA modules operating simultaneously at two different conditions for seven days. Each module had a H_2_O_2_ feedback controller and a PID temperature controller with Module 1 set at 67° C and Module 2 set at 87° C. Values were logged with a Raspberry Pi which was connected to each feedback controller to read the process values. Noise in the H_2_O_2_ readings was largely in part due to the magnetic stir bar.

## Results

The new aRAA system has redesigned heating, automated control of H_2_O_2_ concentration and automated logging of the data. The original RAA design used oil heating with 1000 ml jacketed flask to enable accelerated aging at high temperature. This design was prone to failure due to rapid degradation of oil tubing and eventual ruptures with oil spillage. The use of electric mantles for heating enabled miniaturization of the aRAA setup with a ten-fold decrease in the volume of a reaction flask without change in system performance. Original RAA system does not have temperature login ability, but based on day-to-day observations, it was oscillating within several degrees Celsius. In the new aRAA design, the temperature was 87 ± 1.5° C and 67 ± 4° C for the first and second RAA modules, respectively (Figure 3A).

The main breakthrough of the new aRAA design that makes it simple and easy to use is a feedback loop for automated control of H_2_O_2_ concentration. Chronoamperometric pulsing protocol with 1 sec cathodic hold at - 0.3 V, followed by 2 sec anodic pulse at + 0.7 V, demonstrated stable reading of H_2_O_2_ over a period of time sufficient to perform entire RAA experiments (Figure 3B). The system has not been optimized to remove or minimize electronic noise, which can introduce what appear as sharp spikes in H_2_O_2_ concentration. Most of this noise has been traced to electrical interference from a magnetic stirrer. Since experiments are run on a long time scale (days and weeks) and H_2_O_2_ degradation happens relatively slow (half-life ~ 20 minutes) compared to sampling rate (1 sample every 3 seconds), this interference can be addressed algorithmically in a Python code on Raspberry Pi that determines when there is a glitch and when it is a real drop or spike in H2O2 concentration. This approach allowed us to use readily available Rodeostat without need to optimize detection conditions and by making it easy to use to anybody. Precision of initial version of RAA system on maintaining H_2_O_2_ concentration was a function of operator and typically stayed within 10 mM. The new aRAA design has peak-to-peak H_2_O_2_ concentration within 3 mM (Figure 3B).

Simplicity of the aRAA system and precise control of temperature and H_2_O_2_ concentration enables easy scaling of RAA experiments, with the ability to adjust intensity of aggressive conditions to simulate different degrees of device degradation. To take advantage of this new feature, two types of neural implants were exposed to two different sets of conditions. Blackrock microelectrode arrays were exposed to conditions reported earlier (15 mM H_2_O_2_ at 87° C) (Takmakov et al., 2015), while TDT microelectrode arrays were exposed to milder conditions (15 mM H_2_O_2_ at 67° C). The results obtained for aRAA of Blackrock closely resembles what has been reported earlier including partial cracking and etching of Parylene-C insulation (Figure 4A-D) as well as a drop in electrode impedance over the wide range of frequencies (Figure 4E-F), likely caused by penetration of moisture through the insulation.

**Figure 4.**
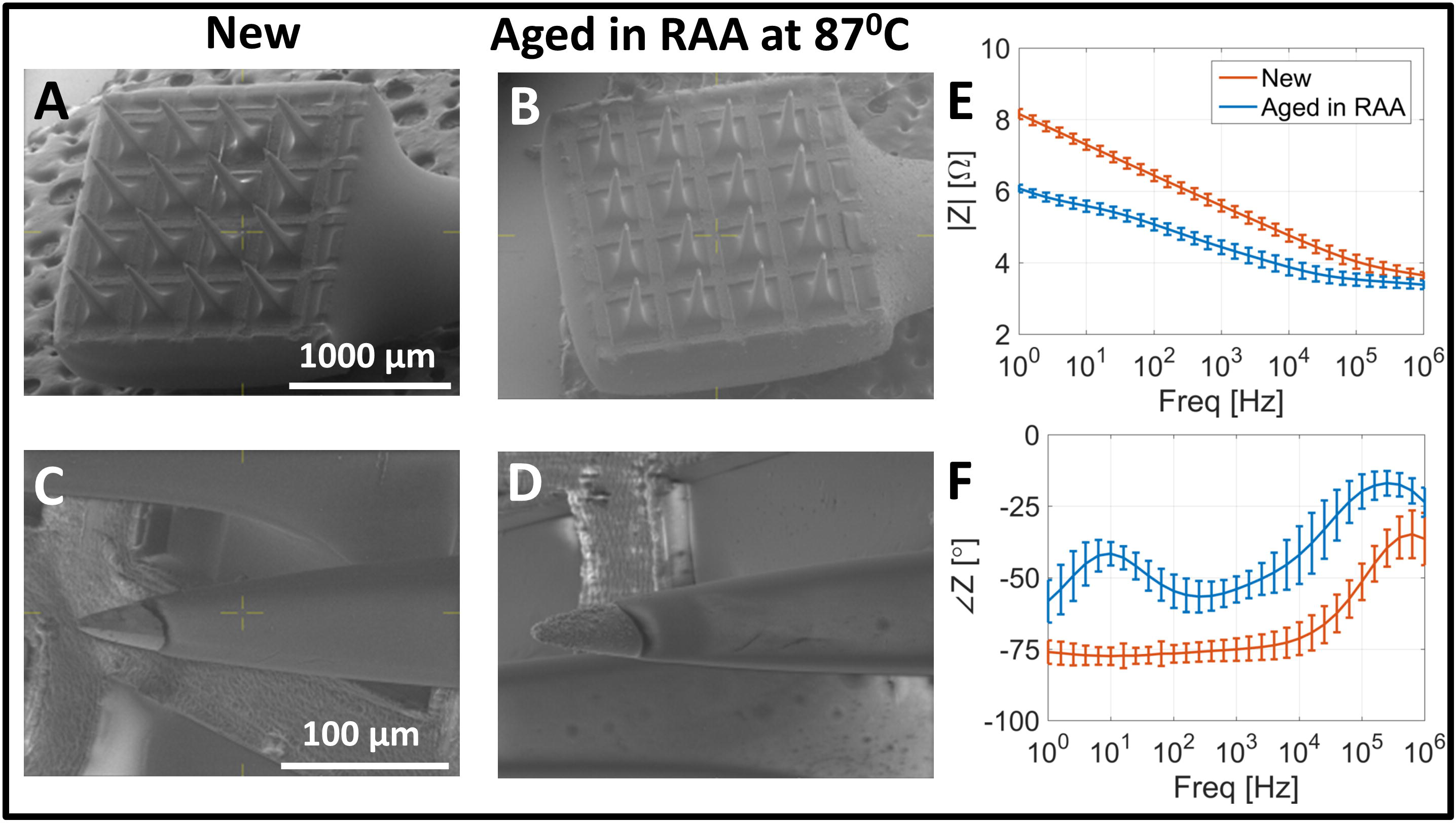
Degradation of Blackrock microelectrodes array in automated RAA system at 87° C. Electron microscopy indicates delamination and etching of Parylene-C insulation (**A** vs **B** and **C** vs **D**). Impedance spectroscopy indicates drop in impedance (**E**) and shift from capacitive into more resistive conduction mode for lower frequencies (**F**).

Examination of TDT MEAs after aRAA at the milder conditions revealed a less extreme degradation pattern than what has been observed earlier (Takmakov et al., 2015) for a higher temperature (87° C). TDT microelectrodes have a tungsten core that is covered with a thin gold layer and insulated with polyimide. Exposure of TDT arrays to RAA at 87° C led to complete dissolution of tungsten core, an almost complete loss of polyimide insulation, and bending and breaking of thin walled gold tubes. However, after aRAA at 67° C, none of the microelectrodes were broken (Figure 5A-B) with cracks and loss of polyimide insulation for some of the electrodes (Figure 5C-F). The impedance data is consistent with earlier RAA experiments with little change after RAA (Figure 5G-H). This suggests that the aRAA experiment with TDT arrays at 67° C appears to be more similar to degradation reported of TDT arrays after in vivo implantation (Prasad and Sanchez, 2012; Prasad et al., 2014), where damage to polyimide insulation was also less severe than what has been observed in original RAA (Takmakov et al., 2015). The ability to tune strength of reactive accelerated aging with better control over experimental parameters (temperature, H_2_O_2_ concentration and time) combined with the ability to replicate the aRAA setup and run multiple experiments in parallel will enable the search for optimal conditions that more accurately match degradation of the implants observed *in vivo*.

**Figure 5.**
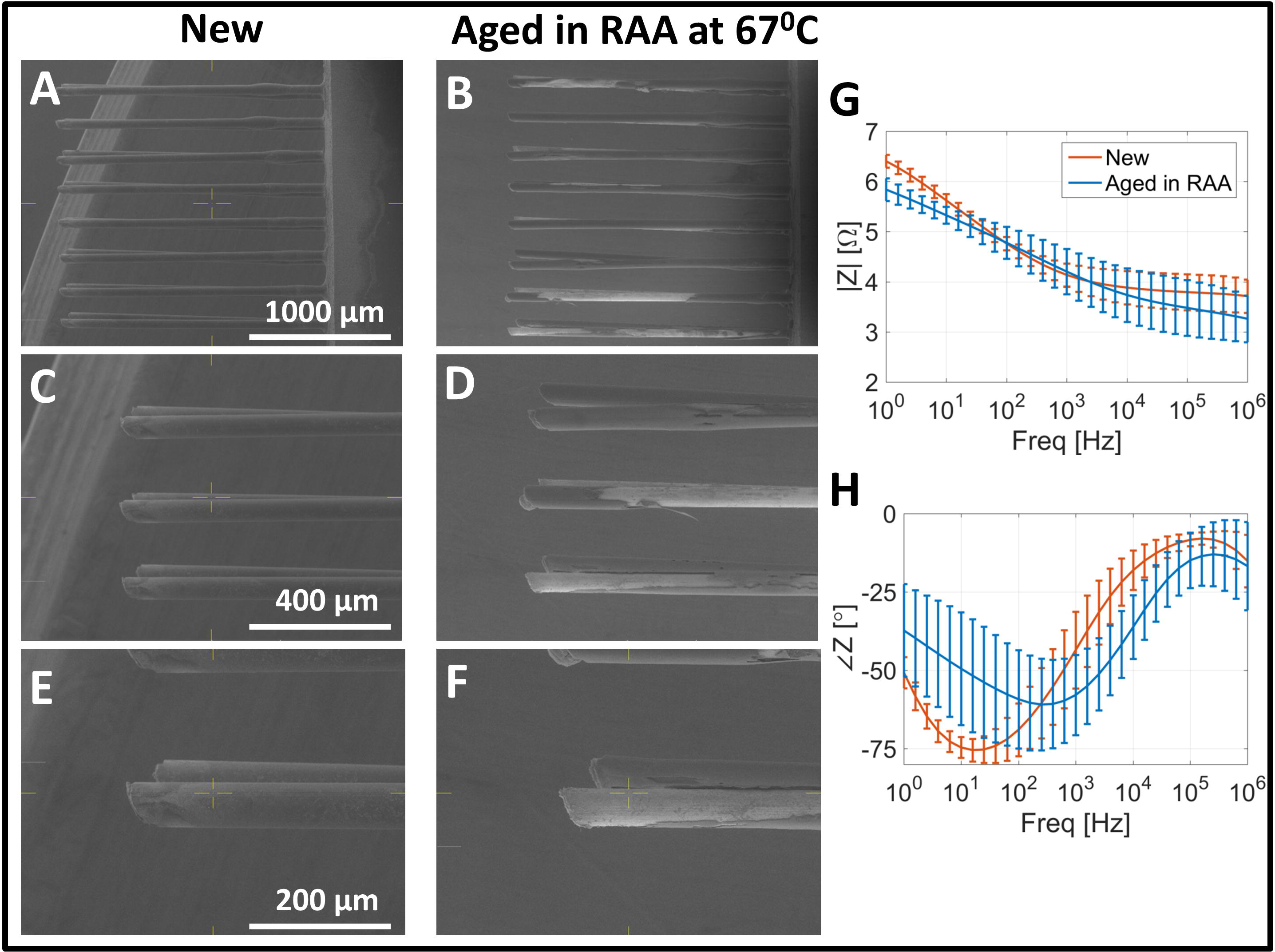
Degradation of TDT microelectrodes array in the automated RAA system at 67° C. Electron microscopy indicates delamination of polyimide insulation and dissolution of tungsten electrode cores (**A** vs **B**, **C** vs **D** and **E** vs **F**), while impedance spectroscopy does not indicate significant change in impedance magnitude (**G**) or phase (**H**).

## Discussion

Automated reactive accelerated aging (aRAA) is a simple setup that can be used for rapid screening of new neural implants to establish potential failure modes without the need for lengthy and expensive animal experiments. The new version of RAA system is fully automated and implements feedback loop to maintain H_2_O_2_ concentration using simple electrochemical detection protocol. The aRAA design employs readily available components and open-source solutions and can be easily replicated on a modest budget. Simplicity of aRAA system and precision in control over temperature and H_2_O_2_ concentration enables easy scaling of RAA experiments and to tune intensity of aggressive conditions to simulate different degree of device degradation.

## Acknowledgements

This work was sponsored by the Defense Advanced Research Projects Agency (DARPA) BTO under the auspices of Dr. Doug Weber through the DARPA Contracts Management Office. Grant/Contract: Inter Agency Agreement with U.S. Food and Drug Administration. We are grateful to Dr. Yong Wu and Dr. Jiwen Zheng of Advanced Characterization Facility (US FDA) for technical assistance with electron microscopy. We thank Dr. Katherine Vorvolakos (US FDA) for review and valuable input to the manuscript.

The views, opinions, and/or findings contained in this article are those of the authors and should not be interpreted as representing the official views or policies of the Department of Defense or the U.S. government. The mention of commercial products, their sources, or their use in connection with material reported herein is not to be construed as either actual or implied endorsement of such products by the Department of Health and Human Services.

